# Naturally occurring ZCCHC3 variants modulate antiretroviral activity in cynomolgus macaques

**DOI:** 10.64898/2026.05.21.726815

**Authors:** Jacob Fadipe, Tomotaka Okamura, Shige H Yoshimura, Akatsuki Saito

**Author notes:** These authors contributed equally (Jacob Fadipe, Tomotaka Okamura).

## Abstract

Many mammalian cells restrict viral replication by utilizing various host restriction factors. We recently demonstrated that CCHC-type zinc-finger-containing protein 3 (ZCCHC3) suppresses human immunodeficiency virus type 1 (HIV-1) replication through multiple mechanisms. We also revealed that single-nucleotide polymorphisms (SNPs) in human ZCCHC3 affect its antiviral function; however, whether similar genetic and functional diversity is present in other species remains unknown.

In this study, we investigated the genetic and functional diversity of ZCCHC3 in cynomolgus macaques, a critical animal model for HIV-1-related research. Sequencing analysis of eight independent ZCCHC3 clones per animal revealed substantial amino acid diversity among cynomolgus macaques. We selected 12 representative variants and examined their antiviral activity against several retroviral vectors derived from HIV-1, simian immunodeficiency virus, feline immunodeficiency virus, and murine leukemia virus. Moreover, using replication-competent HIV-1, we showed that selected cynomolgus macaque ZCCHC3 variants can affect both viral production and viral infectivity. These results suggest that the genetic and functional diversity of ZCCHC3 is not limited to humans and underscore the importance of considering ZCCHC3 variation in cynomolgus macaques when using them as animal models for HIV-1-related research.

## 1. Introduction

Human immunodeficiency virus type 1 (HIV-1) remains a major global health burden [1] despite the widespread use of antiretroviral therapy [2]. Although current treatment regimens efficiently suppress viral replication [3], they do not eliminate the viral reservoir [4], and lifelong therapy is generally required [3,5]. Therefore, a better understanding of the host determinants that regulate HIV-1 replication is essential for elucidating viral pathogenesis and developing new therapeutic strategies [6].

Host restriction factors constitute a critical component of intrinsic antiviral immunity [7]. These factors suppress viral replication at various steps of the viral life cycle and often contribute to species specificity, tissue tropism, and inter-individual variation in viral susceptibility [7,8]. In the context of HIV-1 research, such host factors are particularly important because the development of fully permissive and pathogenic animal models remains challenging [9,10,11]. Non-human primates [12, 13], including cynomolgus macaques [14], have been widely used in studies employing simian immunodeficiency virus (SIV), simian-human immunodeficiency virus (SHIV), and macaque-tropic HIV-1 derivatives [11,12]. However, viral replication and disease progression can vary substantially among individual animals, suggesting that host genetic background may influence experimental outcomes [15].

CCHC-type zinc-finger-containing protein 3 (ZCCHC3) has recently emerged as an antiviral host factor [16,17,18]. We previously demonstrated that human ZCCHC3 suppresses HIV-1 replication through multiple mechanisms [16], including inhibition of viral production and impairment of viral infectivity. We also showed that single-nucleotide polymorphisms (SNPs) in human ZCCHC3 can alter its antiviral activity [16]. These findings raise the possibility that naturally occurring ZCCHC3 polymorphisms may contribute to inter-individual differences in susceptibility to retroviral infection [16]. However, whether ZCCHC3 exhibits similar genetic and functional diversity in non-human primates remains unknown.

In this study, we investigated the genetic and functional diversity of ZCCHC3 in cynomolgus macaques, an important animal model for HIV-1-related research. We identified ZCCHC3 variants in cynomolgus macaques and examined the antiviral activities of selected representative variants against multiple retroviral vectors, including HIV-1-, SIV-, feline immunodeficiency virus (FIV)-, and murine leukemia virus (MLV)-derived vectors, as well as replication-competent HIV-1. Our findings demonstrate that naturally occurring ZCCHC3 variants in cynomolgus macaques modulate antiretroviral activity. These results suggest that the genetic and functional diversity of ZCCHC3 is not limited to humans and should be considered when cynomolgus macaques are used as animal models for HIV-1-related research.

## 2. Materials and methods

### 2.1 Identification of cynomolgus macaque ZCCHC3 variants by cDNA cloning

Blood was collected from 63 cynomolgus macaques housed at the Tsukuba Primate Research Center (TPRC) of the National Institutes of Biomedical Innovation, Health, and Nutrition (NIBN) under anesthesia with ketamine hydrochloride (7.5 mg/kg) and xylazine (3 mg/kg). All procedures were approved by the Animal Care and Use Committee of NIBN (Approval No. DS24-28) and conducted in accordance with NIBN’s animal experimentation regulations and relevant institutional guidelines.

Total RNA was isolated from peripheral blood mononuclear cells (PBMCs) of cynomolgus macaques using the RNeasy Mini Kit (Qiagen, Germany, Cat. No. 74104) according to the manufacturer’s instructions. cDNA was synthesized using an oligo(dT) primer and the Omniscript Reverse Transcription Kit (Qiagen, Cat. No. 205113).

The target gene was amplified by PCR using PrimeSTAR Max DNA Polymerase (TaKaRa, Shiga, Japan, Cat. No. R045A) with the following primers: forward, 5′-TCTCGAGCTCAAGCTTATGGCCACCGGCGGCGGCGCGGAGG-3′; and reverse, 5′-ATCCCCGCGGCCGCGGTACCTTAGTGCCCGGCCACGCCGGTTAG-3′. PCR was performed under the following conditions: initial denaturation at 95°C for 2 min, followed by 30 cycles of denaturation at 95°C for 10 s and annealing/extension at 68°C for 30 s. The amplified products were purified and inserted into the pCMV-HA-N vector (TaKaRa, Cat. No. 635690) using the In-Fusion Snap Assembly Master Mix (TaKaRa, Cat. No. 638947). Eight independent clones per animal were selected, and the resulting constructs were verified by Sanger sequencing performed by a commercial sequencing service (FASMAC).

### 2.2 Plasmids

The following plasmids were obtained through the NIH HIV Reagent Program, Division of AIDS, NIAID, NIH: Human Immunodeficiency Virus 1 (HIV-1), Strain NL4-3 Infectious Molecular Clone (pNL4-3) [19], (Cat. No. ARP-114), contributed by Dr. M. Martin, SIV Packaging Construct (SIV3+, Cat. No. ARP-13456) and SIV LTR Luciferase mCherry Reporter Vector (Cat. No. ARP-13455), both of which were provided by Dr. Tom Hope. The psPAX2-IN/HiBiT and pWPI-Luc2 plasmids were kind gifts from Dr. Kenzo Tokunaga [20]. pCPRDEnv (Cat. No. 1732; http://n2t.net/addgene:1732; RRID: Addgene_1732) was a gift from Dr. Garry Nolan. pMD2.G was a gift from Dr. Didier Trono (Cat. No. 12259; http://n2t.net/addgene:12259; RRID: Addgene_12259). A luciferase-encoding FIV vector, pLionII-luc2 plasmid, and a luciferase-encoding MLV vector, pDON-5 Neo-luc2 plasmid, were described previously [21].

### 2.3 Cell Culture

TZM-bl cells (HRP-8129) were obtained through BEI Resources, NIAID, NIH [22, 23, 24, 25, 26]. TZM-bl cells and Lenti-X 293T cells (TaKaRa, Cat. No. Z2180N) were cultured in Dulbecco’s modified Eagle’s medium (Nacalai Tesque, Kyoto, Japan, Cat. No. 08458-16) supplemented with 10% fetal bovine serum (Cytiva, Marlborough, MA, USA, Cat. No. SH30396) and 1× Penicillin-Streptomycin Mixed Solution (Stabilized) (Nacalai Tesque, Cat. No. 09367-34) at 37°C in a humidified incubator with 5% CO₂. Human CD4+ T cell-derived MT4 cells were cultured in RPMI 1640 medium with L-glutamine (Nacalai Tesque, Cat. No. 30264-56), supplemented with 10% fetal bovine serum and 1× Penicillin-Streptomycin Mixed Solution (Stabilized), at 37°C in a humidified incubator with 5% CO₂.

### 2.4 Production of viruses

To produce an HIV-1-based lentiviral vector, Lenti-X 293T cells were seeded in a 12-well plate (FUJIFILM Wako Pure Chemical, Cat. No. 636-28421) at 2.5 × 10⁵ cells/well and co-transfected with 0.3 µg psPAX2-IN/HiBiT, 0.3 µg pWPI-Luc2, 0.15 µg pMD2.G, and 0.15 µg pCMV-HA-N vectors expressing each selected ZCCHC3 variant. Transfection was performed using TransIT-LT1 Transfection Reagent (TaKaRa, Cat. No. V2304T) in Opti-MEM (Thermo Fisher Scientific, Cat. No. 31985062). To produce an SIV-based lentiviral vector, Lenti-X 293T cells were co-transfected with 0.3 µg pSIV3+, 0.3 µg pSIV-Luc-mCherry, 0.15 µg pMD2.G, and 0.15 µg pCMV-HA-N vectors expressing each selected ZCCHC3 variant. To produce an FIV-based vector, Lenti-X 293T cells were co-transfected with 0.3 µg pCPRDEnv, 0.3 µg pLionII-luc2, 0.15 µg pMD2.G, and 0.15 µg pCMV-HA-N vectors expressing each selected ZCCHC3 variant. To produce an MLV-based retroviral vector, Lenti-X 293T cells were co-transfected with 0.3 µg pGP Vector, 0.3 µg pDON-5 Neo-luc2, 0.15 µg pMD2.G, and 0.15 µg pCMV-HA-N vectors expressing each selected ZCCHC3 variant. Culture supernatants were collected and filtered 2 days after transfection.

To produce replication-competent HIV-1_NL4-3_ viruses, Lenti-X 293T cells were seeded in a 12-well plate at 2.5 × 10⁵ cells/well and co-transfected with 0.75 µg pNL4-3 and 0.25 µg pCMV-HA-N vectors expressing each selected ZCCHC3 variant. To investigate the effect of each ZCCHC3 variant on viral production, we measured the amount of reverse transcriptase in the virus stocks using a SYBR Green PCR-enhanced reverse transcription (SG-PERT) assay as previously described [27].

### 2.5 Western blotting

Lenti-X 293T cells were seeded in a 12-well plate at 2.5 × 10⁵ cells/well and co-transfected with 0.75 µg pNL4-3 and 0.25 µg pCMV-HA-N vectors expressing each selected ZCCHC3 variant. At 2 days post-transfection, the cells were washed and lysed in 1× NuPAGE LDS Sample Buffer (Thermo Fisher Scientific, Cat. No. NP0007) containing 2% (v/v) β-mercaptoethanol and incubated at 70°C for 10 min. The expression of HA-tagged ZCCHC3 variant proteins was assessed using a Simple Western Abby system (ProteinSimple, San Jose, CA, USA) with a 12–230 kDa Separation Module, 8 × 25 capillary cartridge (ProteinSimple, Cat. No. SM-W004A). An anti-HA tag mouse monoclonal antibody (6E2; Cell Signaling Technology, Cat. No. 2367S), RePlex Module (ProteinSimple, Cat. No. RP-001), and Anti-Mouse Detection Module (ProteinSimple, Cat. No. DM-002) were used for detection. The amount of input protein was measured using the Total Protein Detection Module (ProteinSimple, Cat. No. DM-TP01).

### 2.6 Infection assay

MT4 cells were seeded in a 96-well plate (FUJIFILM Wako Pure Chemical, Cat. No. 638-28481) at 3.0 × 10⁴ cells/well. The cells were infected with 10 µL/well of each reporter virus produced in the presence of each selected cynomolgus macaque ZCCHC3 variant.

TZM-bl cells were seeded in a 96-well plate (FUJIFILM Wako Pure Chemical, Cat. No. 635-28511) at 1.0 × 10⁴ cells/well. After overnight culture, the cells were infected with 10 µL/well of each HIV-1_NL4-3_ virus produced in the presence of each selected cynomolgus macaque ZCCHC3 variant.

At 2 days post-infection, the infected cells were lysed using the britelite plus (PerkinElmer, Cat. No. 6066769), and luminescence was measured using a GloMax Explorer Multimode Microplate Reader (Promega).

### 2.7 Statistical analysis

Data are presented as the mean ± SD of six technical replicates from one assay and are representative of at least three independent experiments. Differences in antiviral activity between WT ZCCHC3 and each ZCCHC3 variant were evaluated using one-way analysis of variance (ANOVA). A p-value of ≤0.05 was considered statistically significant. Statistical analyses were performed using GraphPad Prism version 11.0.0 (GraphPad Software, Boston, MA, USA).

## 3. Results

### 3.1 Identification of ZCCHC3 variants in cynomolgus macaques

To identify naturally occurring sequence variations in cynomolgus macaque ZCCHC3, total RNA was extracted from peripheral blood mononuclear cells (PBMCs) of cynomolgus macaques, and the ZCCHC3 coding region was amplified by RT-PCR. The amplified products were cloned into a mammalian expression vector, and eight independent clones per animal were subjected to Sanger sequencing. This analysis identified a wild-type (WT) sequence and multiple ZCCHC3 variants among the cynomolgus macaques examined. In this study, WT refers to a ZCCHC3 amino acid sequence identical to the NCBI reference sequence for *Macaca fascicularis* ZCCHC3 (XP_005568656.1).

Among the 63 cynomolgus macaques analyzed, 25 animals yielded only WT ZCCHC3 sequences among the eight clones examined, whereas the remaining animals yielded one or more ZCCHC3 amino acid variants. Specifically, 19 animals yielded a single amino acid variant, 9 animals yielded multiple distinct variants in different clones, and 10 animals yielded clones carrying multiple amino acid substitutions, together with additional distinct variant sequences in other clones. These findings indicate that cynomolgus macaque ZCCHC3 exhibits substantial sequence diversity at the transcript level. Because comprehensive functional characterization of all detected variants was beyond the scope of this study, we selected 12 representative amino acid variants for further analysis based on their occurrence in the sequenced clones and their positions within predicted functional regions of ZCCHC3.

The selected variants included S72G/A118S, G76S/A131T, G95R/A131T, A118S, A118S/L396P, A120T, A131T, V263M, Q304*, E306G, R351G, and L373P. The amino acid sequences of these cynomolgus macaque ZCCHC3 variants were aligned with human (NP_149080.2) and rhesus macaque ZCCHC3 (XP_001111701.1) sequences (**Figure 1A**). Mapping these substitutions onto the predicted domain structure of ZCCHC3 showed that several variants, including S72G/A118S, G76S/A131T, G95R/A131T, A118S, A118S/L396P, A120T, and A131T, were located in the N-terminal disordered region (**Figure 1B**). V263M, Q304*, and E306G were located within the central fold domain. Notably, Q304* introduced a premature stop codon, resulting in truncation of the C-terminal region containing the zinc-finger motifs. The remaining variants, R351G and L373P, were located in the C-terminal region containing the zinc-finger motifs (**Figure 1B**). These results indicate that cynomolgus macaque ZCCHC3 exhibits amino acid diversity across multiple regions of the protein, and that the selected variants provide a useful panel for assessing the functional consequences of naturally occurring ZCCHC3 variation.

**Figure 1.**
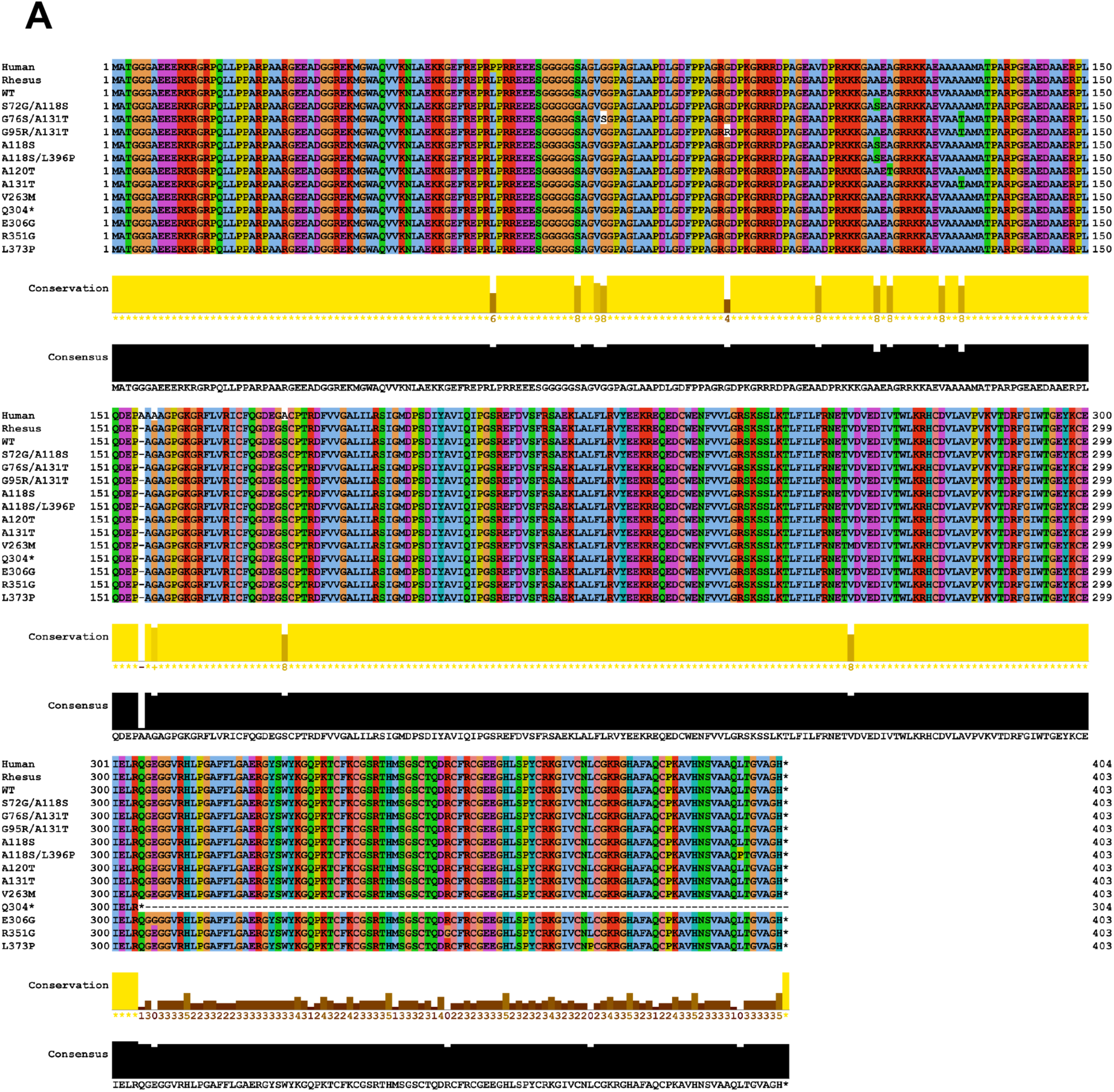
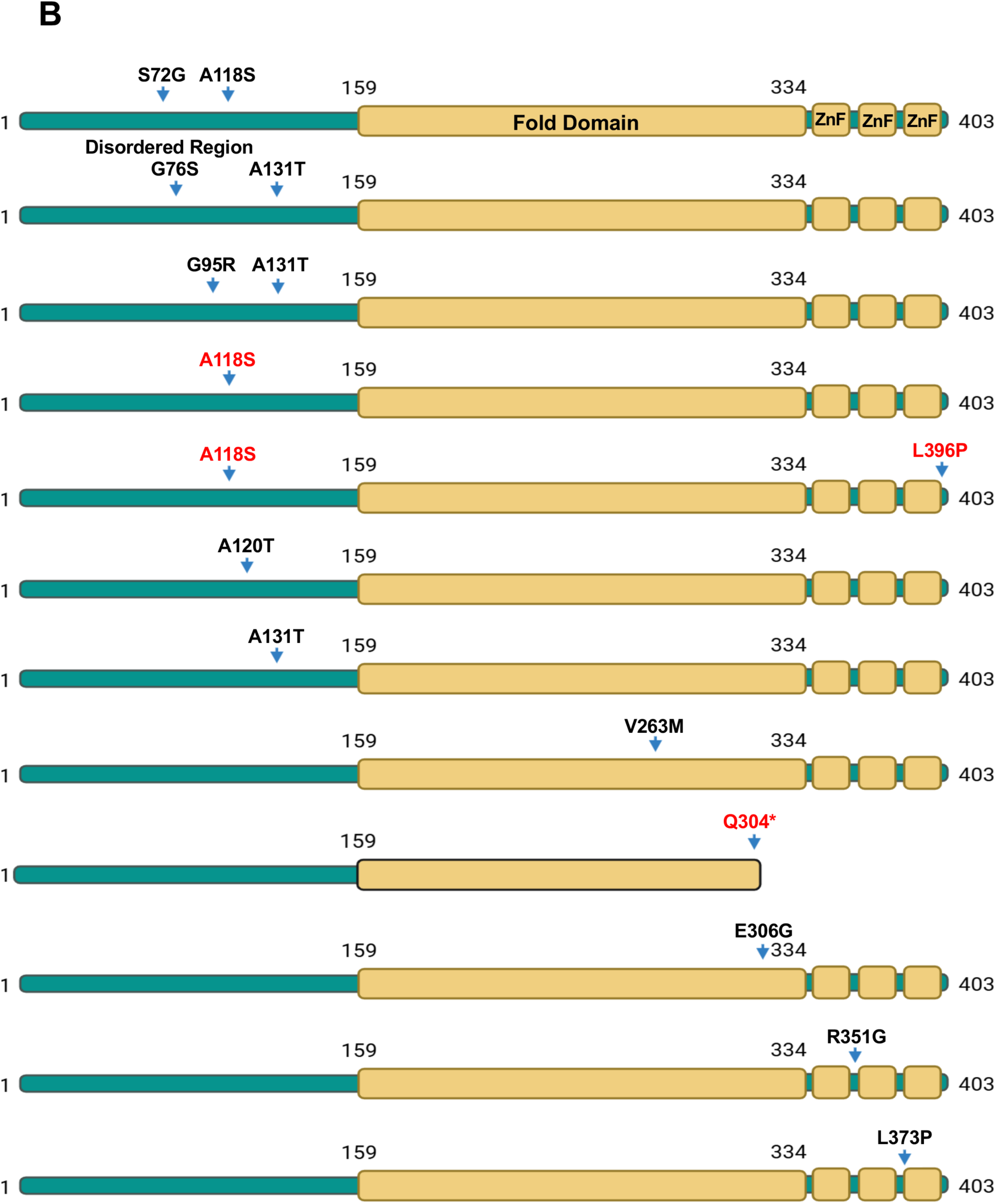
Identification and domain mapping of ZCCHC3 variants selected for functional analysis. (A) Amino acid sequence alignment of human, rhesus macaque, and selected cynomolgus macaque ZCCHC3 variants. (B) Schematic representation of the ZCCHC3 protein showing the positions of amino acid substitutions selected for functional analysis in this study.

### 3.2 Expression of cynomolgus macaque ZCCHC3 variants

We next examined the expression of HA-tagged cynomolgus macaque ZCCHC3 variants selected for functional analysis using the Simple Western Abby system. Most ZCCHC3 variants were detected at an apparent molecular weight of approximately 45–55 kDa (**Figure 2A**). The Q304* variant migrated at a lower molecular weight than the other variants, consistent with the predicted C-terminal truncation caused by the premature stop codon. Total protein detection was used to confirm sample loading (**Figure 2B**). These results confirmed that the ZCCHC3 variants used in this study were expressed at detectable levels in transfected cells.

**Figure 2.**
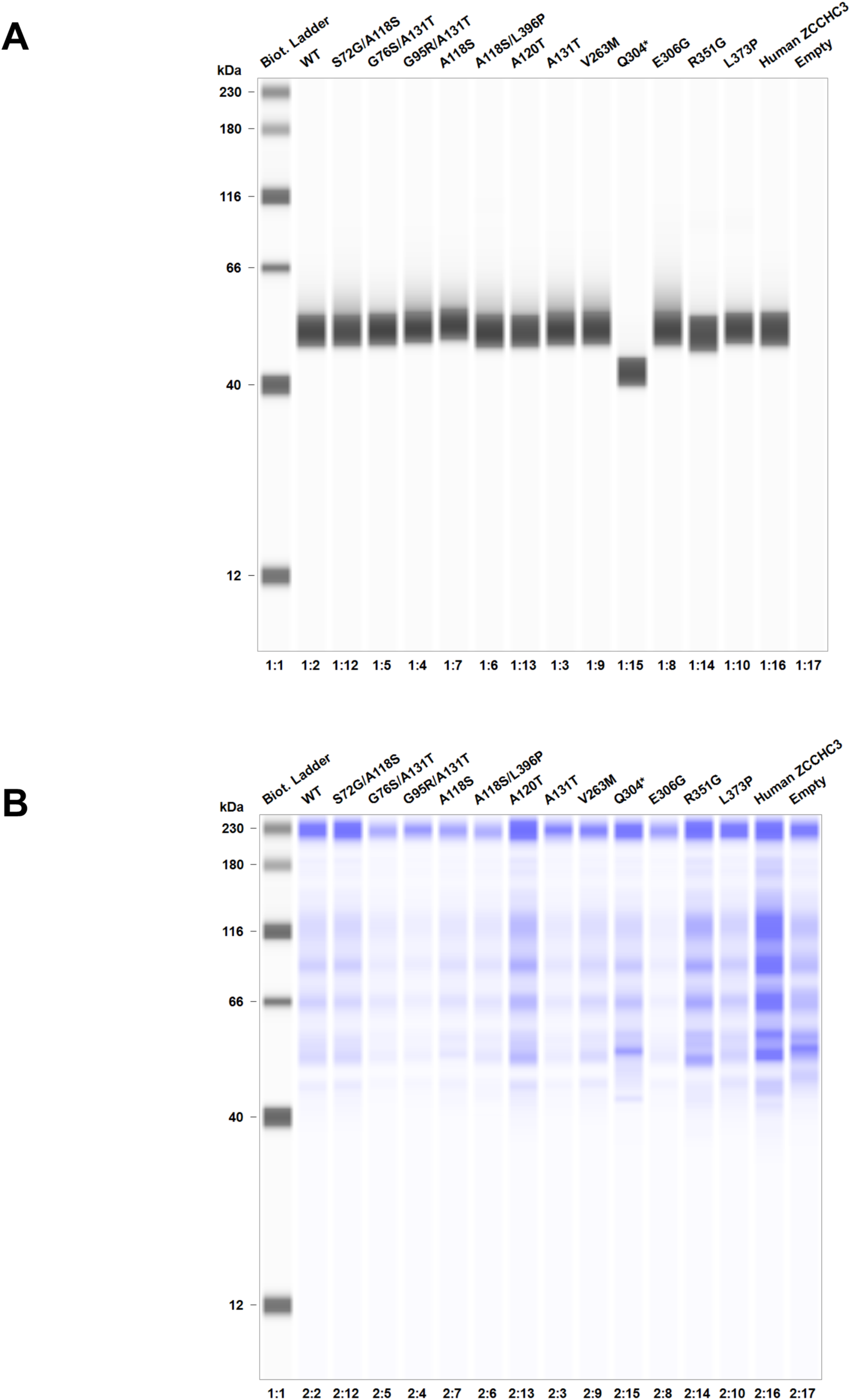
Expression of selected cynomolgus macaque ZCCHC3 variants. (A) Lenti-X 293T cells were co-transfected with the pNL4-3 plasmid and pCMV-HA-N vectors encoding each cynomolgus macaque ZCCHC3 variant. Human ZCCHC3 indicates full-length human ZCCHC3 and was included as a reference control. At 2 days post-transfection, cells were lysed, and the expression of HA-tagged selected cynomolgus macaque ZCCHC3 variants was analyzed using the Simple Western Abby system. (B) Total protein levels were measured as a loading control.

### 3.3 Differential antiviral activity of selected cynomolgus macaque ZCCHC3 variants against HIV-1 and retroviral vectors

To assess the functional impact of the selected ZCCHC3 variants, we first examined the antiviral activity of cynomolgus macaque ZCCHC3 variants against HIV-1-, SIV-, FIV-, and MLV-derived reporter vectors. MT4 cells were infected with reporter viruses produced in the presence of each ZCCHC3 variant, and infection levels were quantified using a luciferase assay. In this assay, lower relative luciferase activity indicates stronger antiviral activity.

Most cynomolgus macaque ZCCHC3 variants showed antiviral activity comparable to that of WT ZCCHC3, although the magnitude of restriction varied depending on the viral vector tested (**Figures 3A, 3B, 3C, and 3D**). Among the selected variants examined, Q304* consistently showed markedly reduced antiviral activity, as indicated by significantly increased relative luciferase activity compared with WT ZCCHC3. The activity of Q304* was close to that of the empty vector control in multiple assays, suggesting that truncation of the C-terminal region containing the zinc-finger motifs severely impairs the antiviral function of ZCCHC3. In contrast, several variants showed virus-dependent effects. For example, A118S/L396P showed significantly lower relative luciferase activity in the MLV-based vector assay, suggesting enhanced antiviral activity in this context (**Figure 3D**). Other variants, such as R351G and L373P, showed apparent changes in relative luciferase activity in some assays, but these differences did not reach statistical significance. Similarly, A120T tended to show reduced relative luciferase activity in the FIV- and MLV-based vector assays but not consistently across all viral vectors.

**Figure 3.**
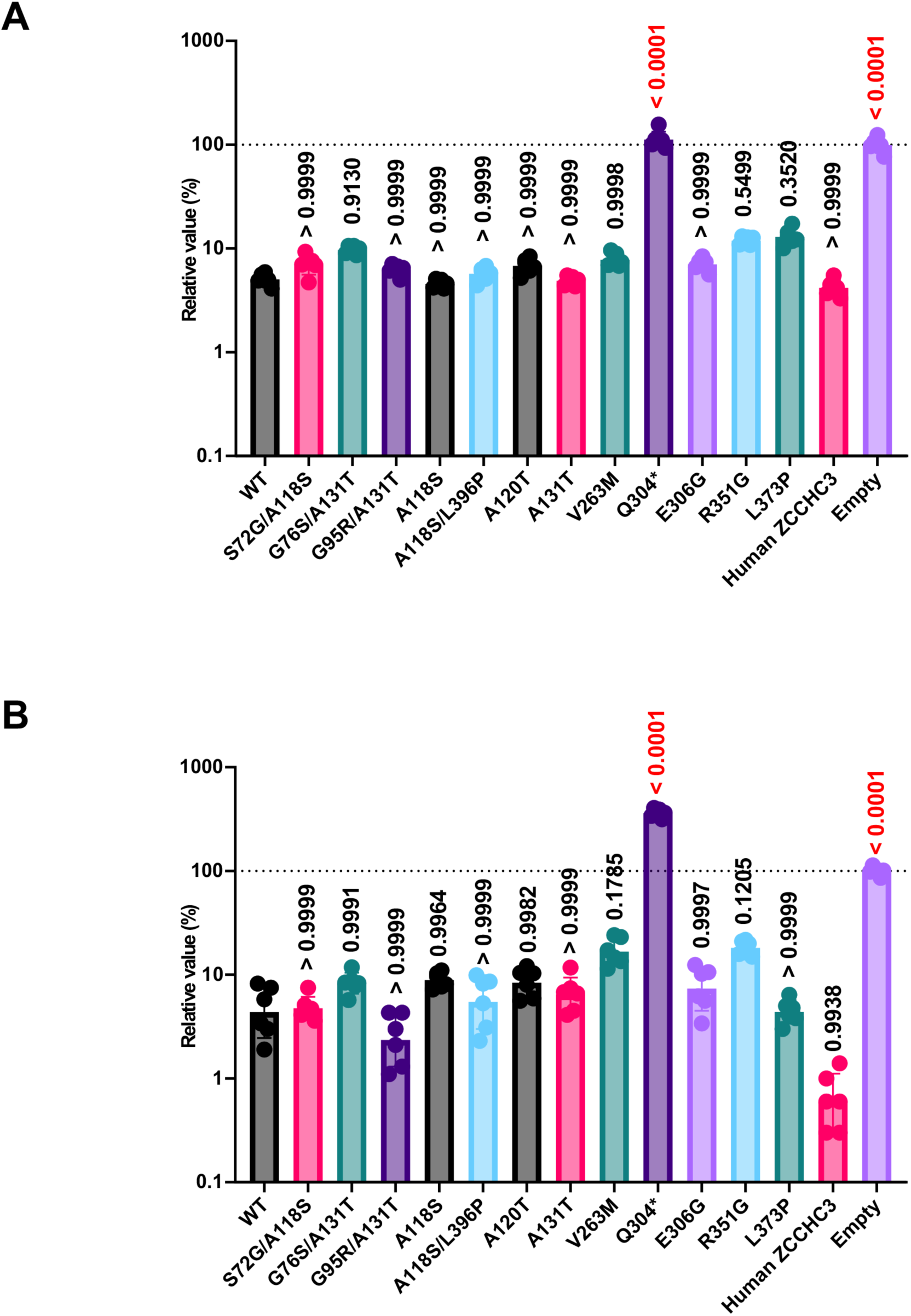
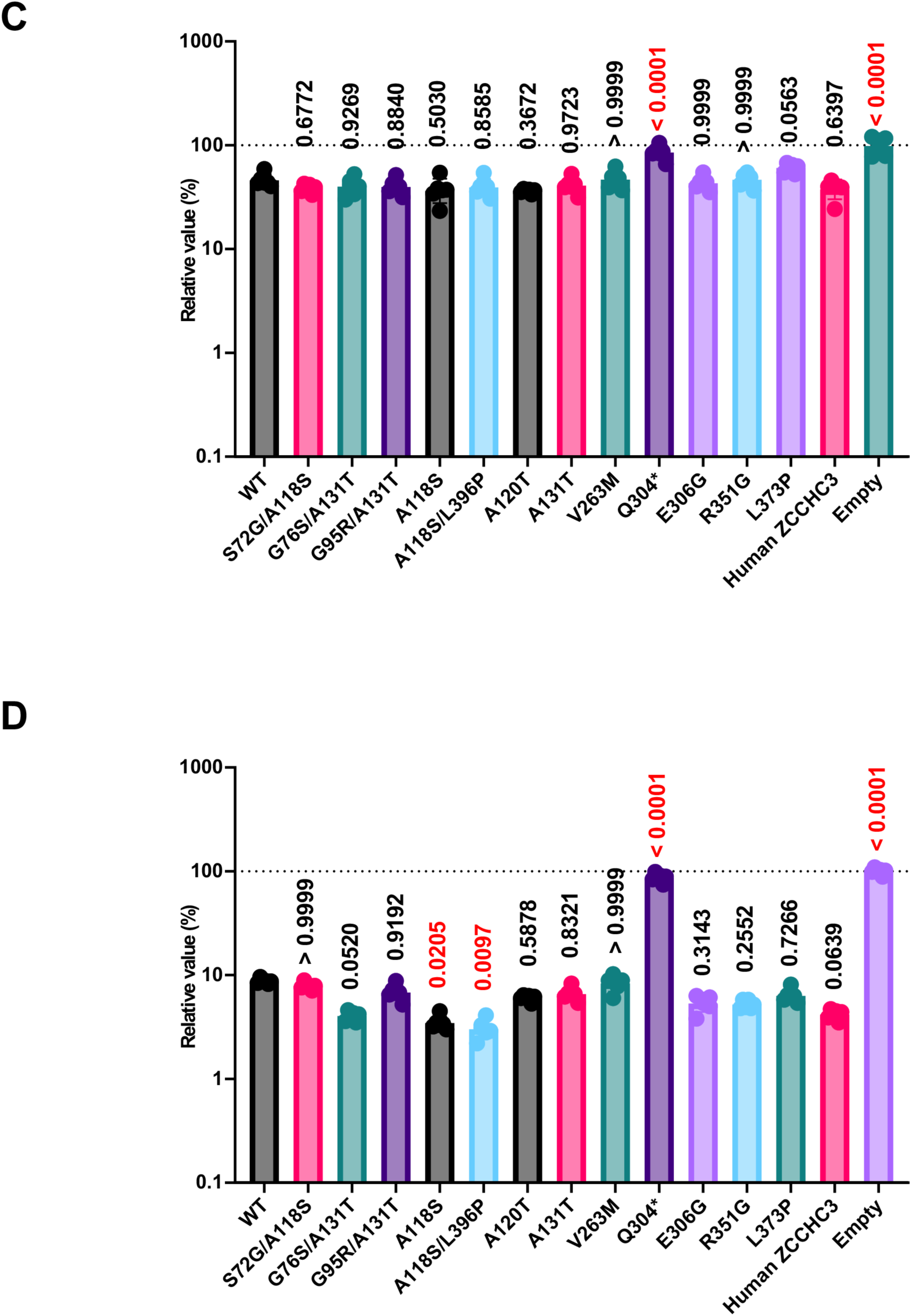
Effects of selected cynomolgus macaque ZCCHC3 variants on retroviral vector infection. MT4 cells were infected with 10 µL/well of each reporter virus produced in the presence of each selected cynomolgus macaque ZCCHC3 variant. Human ZCCHC3 indicates full-length human ZCCHC3 and was included as a reference control. Luciferase activity was measured 2 days after infection. Relative values were calculated by normalizing luciferase activity to that of the empty vector control. Data are presented as the mean ± SD of six technical replicates from one representative assay and are representative of three independent experiments. Statistical significance was analyzed by one-way ANOVA; p < 0.05 was considered statistically significant. (A) HIV-1-based vector. (B) SIV-based vector. (C) FIV-based vector. (D) MLV-based vector.

We next examined whether the selected naturally occurring ZCCHC3 variants influence antiviral activity against replication-competent HIV-1_NL4-3_. Consistent with the results obtained using reporter vectors, most cynomolgus macaque ZCCHC3 variants showed strong antiviral activity against HIV-1_NL4-3_, whereas several variants, most notably Q304*, showed reduced antiviral activity (**Figure 4A**). To further investigate the mechanism underlying this antiviral effect, we quantified virion-associated reverse transcriptase activity using the SG-PERT assay. Most ZCCHC3 variants reduced viral production, whereas Q304* showed little or no inhibitory effect on viral production (**Figure 4B**).

**Figure 4.**
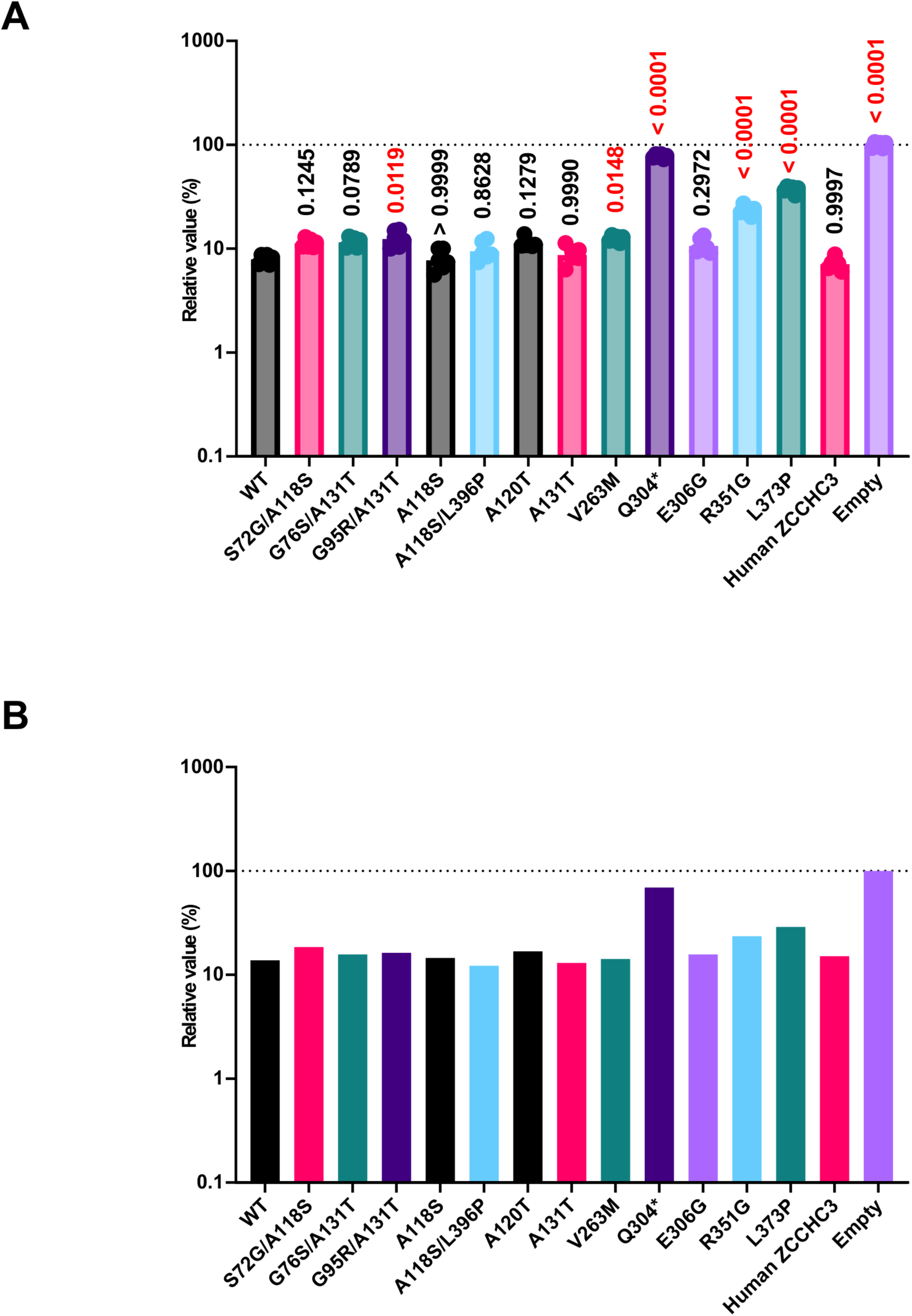
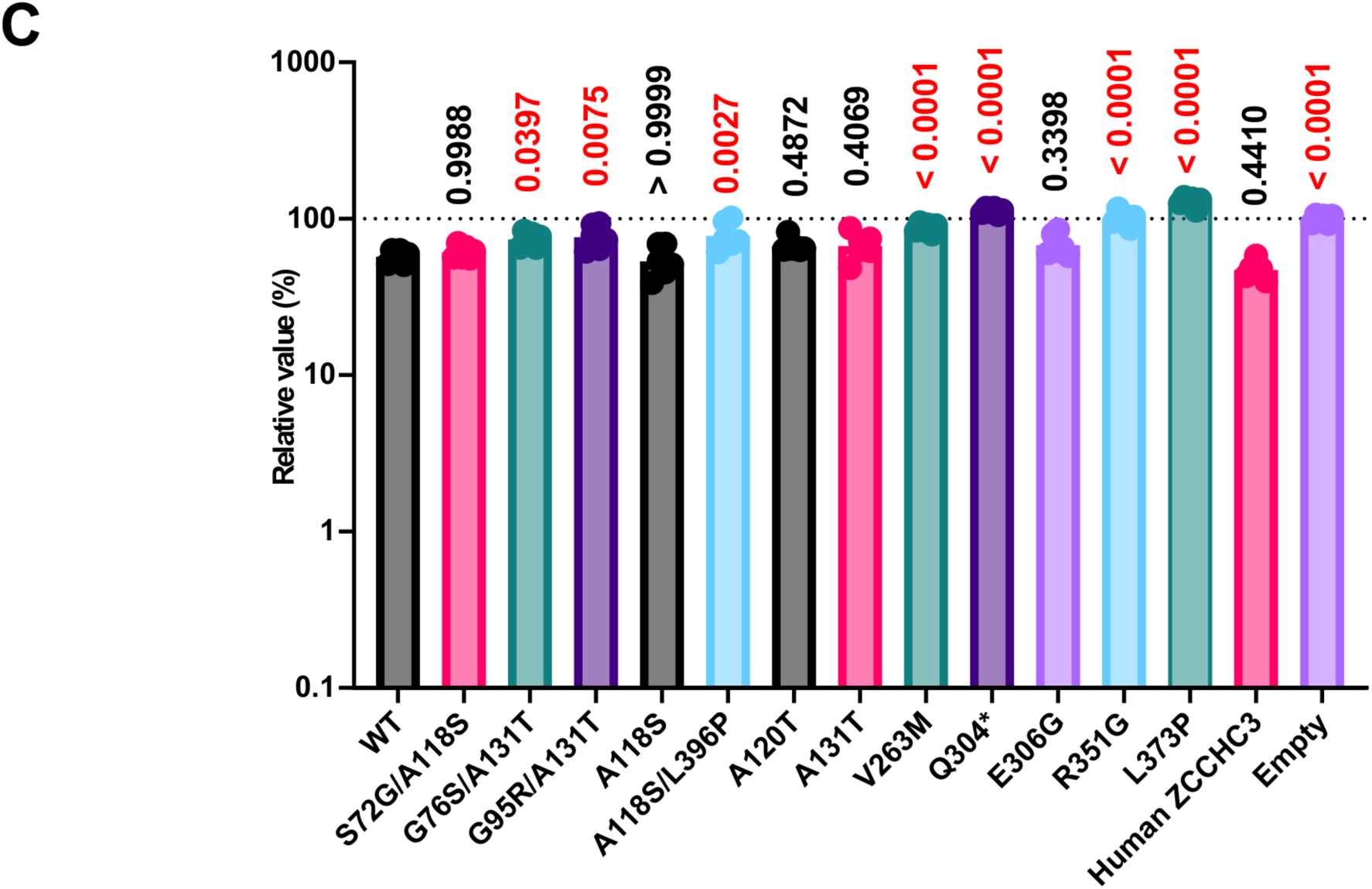
Effects of selected cynomolgus macaque ZCCHC3 variants on replication-competent HIV-1. (A) TZM-bl cells were infected with 10 µL/well of HIV-1_NL4-3_ viruses produced in the presence of each selected cynomolgus macaque ZCCHC3 variant. Human ZCCHC3 indicates full-length human ZCCHC3 and was included as a reference control. Luciferase activity was measured 2 days after infection. Relative values were calculated by normalizing luciferase activity to that of the empty vector control. (B) Viral production from Lenti-X 293T cells in the presence of each selected cynomolgus macaque ZCCHC3 variant was quantified using the SG-PERT assay. (C) Relative infectivity was calculated by dividing the raw luciferase values shown in panel A by the viral production values shown in panel B to normalize for virus input. Data are presented as the mean ± SD of six technical replicates from one representative assay and are representative of three independent experiments. Statistical significance was analyzed by one-way ANOVA; p < 0.05 was considered statistically significant.

Finally, we calculated relative infectivity by normalizing luciferase activity to virion-associated reverse transcriptase activity. This analysis showed that most cynomolgus macaque ZCCHC3 variants reduced relative infectivity, whereas R351G, L373P, and Q304* showed weaker or minimal effects on relative infectivity (**Figure 4C**). These results suggest that cynomolgus macaque ZCCHC3 can suppress replication-competent HIV-1 through effects on both viral production and viral infectivity, and that some naturally occurring variants impair one or both of these antiviral activities.

Together, these results demonstrate that selected naturally occurring ZCCHC3 variants in cynomolgus macaques can influence antiviral activity. In particular, the loss of antiviral activity observed for Q304* highlights the importance of the C-terminal region, including the zinc-finger motifs, for ZCCHC3-mediated restriction of retroviral infection.

## 4. Discussion

In this study, we investigated the genetic and functional diversity of ZCCHC3 in cynomolgus macaques and examined whether naturally occurring ZCCHC3 variants can affect antiretroviral activity. Sequencing analysis of ZCCHC3 transcripts from cynomolgus macaques revealed substantial amino acid diversity, including animals from which multiple distinct ZCCHC3 sequences were obtained. Because comprehensive functional characterization of all detected variants was beyond the scope of this study, we selected 12 representative variants distributed across different regions of ZCCHC3 and evaluated their antiviral effects against HIV-1-, SIV-, FIV-, and MLV-derived reporter vectors, as well as replication-competent HIV-1_NL4-3_.

The selected variants were distributed across the N-terminal disordered region, the central fold domain, and the C-terminal zinc-finger-containing region (**Figure 1**). Most selected variants retained antiviral activity comparable to that of WT ZCCHC3, whereas Q304* showed a marked loss of antiviral activity (**Figures 3 and 4**). These findings indicate that cynomolgus macaque ZCCHC3 is genetically diverse and that at least some naturally occurring variants can substantially affect its antiviral function.

Sequencing of eight independent ZCCHC3 clones per animal revealed that the pattern of ZCCHC3 variation in cynomolgus macaques was more complex than a simple classification into WT and single-variant alleles. Some animals yielded multiple distinct ZCCHC3 sequences from independent clones, suggesting the presence of complex allelic variation, transcript diversity, or other sources of sequence heterogeneity. Therefore, the functional analyses performed in this study should be interpreted as an initial characterization of selected representative variants rather than a comprehensive assessment of all ZCCHC3 variants present in cynomolgus macaques.

The complex sequence diversity observed in cynomolgus macaque ZCCHC3 may not be unique to this species. In our previous study, human ZCCHC3 was also found to contain more than 2,000 SNPs, and selected human ZCCHC3 variants affected antiviral activity [16]. These findings, together with the present results, suggest that ZCCHC3 may represent a genetically variable antiviral host factor in primates. Nevertheless, the functional significance of most ZCCHC3 variants remains unclear, and broader genotype–function analyses will be required to determine how ZCCHC3 diversity contributes to inter-individual differences in retroviral susceptibility.

The most notable variant identified in this study was Q304*. This variant introduces a premature stop codon and is predicted to delete the C-terminal region of ZCCHC3, including the zinc-finger motifs (**Figure 1B**). Consistent with this prediction, Q304* migrated at a lower molecular weight than the other variants in the Simple Western analysis (**Figure 2A**). Functionally, Q304* showed markedly increased reporter virus infection levels, close to those observed with the empty vector control in multiple assays (**Figure 3**). Furthermore, in the replication-competent HIV-1_NL4-3_ assay, Q304* showed a markedly reduced ability to suppress viral production and relative infectivity (**Figure 4**). Given our previous finding that the zinc-finger motifs mediate binding to Gag nucleocapsid and are required for the suppression of viral genome incorporation into virions [16], the loss of antiviral activity in Q304* strongly supports the importance of the C-terminal zinc-finger-containing region for ZCCHC3-mediated restriction.

The results obtained using replication-competent HIV-1 further support the idea that cynomolgus macaque ZCCHC3 can suppress HIV-1 through multiple steps. Most cynomolgus macaque ZCCHC3 variants reduced HIV-1_NL4-3_ infectivity in TZM-bl cells (**Figure 4A**) and also reduced virion-associated reverse transcriptase activity, indicating inhibition of viral production (**Figure 4B**). After normalization of luciferase activity to virion-associated reverse transcriptase activity, most variants still reduced relative infectivity, suggesting that they affect not only viral production but also the infectivity of released virions (**Figure 4C**). This pattern is consistent with our previous finding that human ZCCHC3 suppresses both viral production and viral infectivity by interfering with viral RNA genome incorporation.

The functional phenotypes of variants located in or near the C-terminal region also support this interpretation. The A118S/L396P variant showed enhanced antiviral activity in the MLV-based vector assay (**Figure 3D**). In the replication-competent HIV-1 assay, however, some C-terminal variants, including R351G and L373P, showed weaker effects on relative infectivity after normalization of virus input (**Figure 4C**). Although the effects of these variants were not identical across all assays, their virus- or assay-dependent phenotypes suggest that amino acid substitutions in or near the zinc-finger-containing region may influence the efficiency of ZCCHC3 incorporation into viral particles, its interaction with retroviral Gag proteins, or its ability to interfere with viral genome packaging. Further biochemical studies will be required to determine whether these cynomolgus macaque variants alter GagNC binding, virion incorporation, or viral RNA recruitment.

In contrast, most variants located in the N-terminal disordered region, including S72G/A118S, G76S/A131T, G95R/A131T, A118S, A120T, and A131T, retained antiviral activity broadly comparable to that of WT ZCCHC3 (**Figures 1B and 3**). This suggests that the N-terminal disordered region may tolerate certain naturally occurring amino acid substitutions without a major loss of antiretroviral function, supporting our previous finding that the N-terminal disordered region of human ZCCHC3 is dispensable for antiviral activity [16].

The virus-dependent differences observed among HIV-1-, SIV-, FIV-, and MLV-derived vectors are also consistent with the dual mechanism proposed in our previous study. ZCCHC3 can act through both interaction with GagNC and recognition of viral LTR sequences. Therefore, differences in GagNC structure or viral RNA sequence may influence the sensitivity of each retroviral vector to ZCCHC3-mediated restriction. In the present study, Q304* consistently lost antiviral activity across the reporter vector assays, whereas several other variants showed vector-dependent effects (**Figure 3A–D**). In addition, the replication-competent HIV-1 assay revealed that some variants can differentially affect viral production and relative infectivity (**Figure 4A–C**). These findings suggest that the C-terminal zinc-finger region is broadly important for antiviral function, while the contribution of individual amino acid substitutions may vary depending on the retroviral context and the step of the viral life cycle being evaluated.

These findings have practical implications for non-human primate studies. Genetic variation in host restriction factors can influence the outcome of infection experiments, especially when experimental systems rely on retroviral vectors, chimeric viruses, or replication-competent HIV-1 derivatives. The present study suggests that ZCCHC3 variation may be one of the host genetic factors contributing to inter-individual variability in cynomolgus macaques. In particular, animals carrying loss-of-function or hypomorphic ZCCHC3 variants may exhibit altered susceptibility to retroviral infection. Therefore, ZCCHC3 variant analysis may be useful when designing or interpreting HIV-1-related experiments using cynomolgus macaques. Furthermore, it is important to investigate the genetic and functional diversity of ZCCHC3 in rhesus macaques and pig-tailed macaques, both of which are also important animal models for HIV-1-related research.

There are several limitations to this study. First, although we identified functional differences among cynomolgus macaque ZCCHC3 variants (**Figures 3 and 4**), the underlying mechanisms were not directly examined. In particular, it remains unclear whether the observed phenotypes are caused by altered GagNC binding, impaired RNA binding, changes in P-body localization, altered virion incorporation, or differences in protein expression and stability. Second, although this study included both reporter vector systems and replication-competent HIV-1_NL4-3_, these *in vitro* systems do not fully recapitulate viral replication and pathogenesis *in vivo*. Third, although the results suggest that Q304* disrupts the zinc-finger-dependent antiviral function of ZCCHC3 (**Figures 1B, 3, and 4**), direct biochemical assays will be required to confirm whether this variant fails to bind GagNC, to recruit viral RNA, or to be incorporated into virions. Fourth, although eight independent ZCCHC3 clones per animal were analyzed, this study functionally characterized only a selected panel of 12 variants. Several animals yielded multiple distinct ZCCHC3 sequences from independent clones, and the biological basis of this sequence diversity remains unresolved. Further studies using genomic DNA-based genotyping, allele-specific sequencing, and long-read sequencing will be required to determine whether these sequences reflect allelic variation, transcript-level diversity, paralogous amplification, or technical artifacts, and to clarify the full spectrum of ZCCHC3 diversity in cynomolgus macaques. The presence of extensive ZCCHC3 variation in both humans and cynomolgus macaques highlights the importance of considering ZCCHC3 variation when interpreting retroviral restriction phenotypes across primate species.

In conclusion, this study demonstrates that cynomolgus macaque ZCCHC3 exhibits genetic and functional diversity. Most selected variants retained antiretroviral activity, whereas Q304* showed a marked loss of function, highlighting the importance of the C-terminal zinc-finger-containing region (**Figures 1–4**). Importantly, analyses using replication-competent HIV-1_NL4-3_ further showed that cynomolgus macaque ZCCHC3 variants can affect both viral production and relative infectivity (**Figure 4**). Together with our previous demonstration that human ZCCHC3 restricts HIV-1 through a dual mechanism involving GagNC binding and viral RNA sequestration, the present findings suggest that ZCCHC3-mediated restriction is conserved across primates but can be modulated by naturally occurring polymorphisms. These results underscore the importance of considering ZCCHC3 variation in cynomolgus macaque models of HIV-1-related research.

## Acknowledgments

The authors thank Ms. Nanami X Kato, Ms. Yuri V Fukushima, Ms. Tomoko Nishiuchi, Dr. Asami Oguro-Ando, and the staff of CADIC, University of Miyazaki, for their assistance. We also thank the members and veterinary staff of HAMRI Co., Ltd. for their technical expertise and assistance with animal care. This study was supported by the Frontier Science Research Center, University of Miyazaki.

The following reagents were obtained from BEI Resources, NIAID, NIH: Human Immunodeficiency Virus 1 (HIV-1), Strain NL4-3 Infectious Molecular Clone (pNL4-3), ARP-114, contributed by Dr. M. Martin, TZM-bl Cells, HRP-8129. The following reagents were obtained through the NIH HIV Reagent Program, Division of AIDS, NIAID, NIH: SIV Packaging Construct (SIV3 +), ARP-13456, and SIV LTR Luciferase mCherry Reporter Vector, ARP-13455, both of which were contributed by Dr. Tom Hope. The following reagents were obtained through the Addgene: pCPRDEnv was a gift from Garry Nolan. pMD2.G was a gift from Dr. Didier Trono. The following plasmids were kind gifts from Dr. Kenzo Tokunaga: psPAX2-IN/HiBiT and pWPI-Luc2.

## Ethics declarations

All animal procedures were approved by the Animal Care and Use Committee of the National Institutes of Biomedical Innovation, Health, and Nutrition (NIBN) (Approval No. DS24-28) and were conducted in accordance with NIBN animal experimentation regulations and relevant institutional guidelines.

